# Expert and Novice Visuomotor Strategies in a Sequential Football Task Reflect Distinct Generative Models

**DOI:** 10.64898/2026.10.06.756964

**Authors:** Guillaume Hacques, Pierpaolo Iodice, Aurélien Dehorne, Amine Chaigneau, Giovanni Pezzulo

## Abstract

In visuomotor sequential tasks, chaining actions smoothly requires tight coordination between sensory and motor systems to deal with competing demands, such as reducing uncertainty about future opportunities for action while controlling ongoing ones. In a football-based decision-making task, experts and novices used different strategies to overcome these competing demands, demonstrating a proactive and a sequential visuomotor strategy, respectively. We propose a computational explanation of expertise-related difference in policy selection, grounded in the active inference framework. We simulated the selection of sequence of actions using a partially observable Markov decision process (POMDP) model, and we manipulated three parameters: the reward structure, the prior beliefs about state transition, and the sensory precision. Simulations showed that selection of the proactive strategy relied on strong prior beliefs, high sensory precision and high valuation of fast passing. Conversely, the sequential policy was selected when rapid performance was not highly valued, or under noisy prior beliefs, or lower level of sensory precision. Decreasing sensory precision and adding noise in prior beliefs also decreased decision accuracy. These results suggest that policy selection leading to novice or expert visuomotor strategy reflects differences in the underlying generative model, particularly in how agents value different aspects of the task.

## Background

Achieving a goal rarely requires a single motor action. In everyday life, making a sandwich or a tea involves performing a sequence of subtasks nested within one another (Hayhoe, 2000; Land, 2006; Land et al., 1999; Land & Hayhoe, 2001). For instance, Land et al. (1999) counted that 40 to 50 separate actions were necessary for making a cup of tea. Each of these actions underlies achievement of subtasks contributing to the overall task-goal. These subtasks (or task modules) require sensory information to reduce uncertainty and adequately perform the required action (e.g., checking the water level in the kettle while filling it or locating the cup handle relative to the hand in order to grasp it). However, each subtask imposes its own informational demands that must be met at the right time, requiring tight coordination of the sensory and motor systems to deal with competing demands of the subtasks and to smoothly chain actions together.

Considering behaviors as sequence of sensorimotor decisions provides a useful framework for understanding how sensory systems (in particular the visual system) support action choices (Cisek & Kalaska, 2010; Gordon et al., 2021; Hayhoe, 2017; Pezzulo & Cisek, 2016; Priorelli et al., 2024; Wolpert & Landy, 2012). Within this framework, decisions about where to look and what to do are driven by the need to resolve the uncertainty of the world and to maximize expected reward, and may also be biased by the new opportunities for action that they make available, as proposed by the affordance competition hypothesis (Cisek, 2007; Cisek & Kalaska, 2010). For example, gaze allocation in a virtual driving task was shown to be clearly affected by the reliability of the sources of information and by the relative emphasis on one subtask over another through task instructions (i.e., maintaining a constant speed or following a car at a constant distance) (Sullivan et al., 2012). This experimental manipulation of instructions was thought to increase intrinsic reward value of one of the subtasks, giving it priority over the other. Specifically, drivers orient their gaze to reduce uncertainty only when the source of that uncertainty was relevant to the prioritized subtask. This gaze pattern was successfully approximated by a softmax barrier model, in which gaze allocation (toward the speedometer or the leading car) depended solely on subtask priority and uncertainty (Johnson et al., 2014). In this model, the state estimate for a subtask drifted over time from the true state as the agent spend time allocating their gaze toward irrelevant objects to that subtask. This growing deviation and uncertainty about the true state could only be reduced by reorienting their gaze toward a relevant location for this subtask. Similarly, task-dependent gaze allocation was further confirmed in a virtual navigation task where participants had to pick up some objects or avoid others or combine both tasks (Rothkopf et al., 2016). This study additionally showed that task goals exerted a greater influence on gaze behavior than object saliency. Together, these studies highlight the importance of task goals for gaze dynamics and for how the visual system manages competing sources of information. However, in these tasks, subtasks were performed in parallel and continuously, whereas the challenge in sequential tasks is to perform subtasks one after another, introducing competition between completing the ongoing subtask and obtaining information for the next one.

A growing body of theoretical work frames perception and action within a “predictive brain” hypothesis, according to which the brain maintains internal generative models that predict incoming sensory information (Hayhoe, 2017; Knill & Pouget, 2004; Parr et al., 2022; Rao & Ballard, 1999). Within this framework, perception corresponds to inferring the hidden causes of sensory input by minimizing prediction errors, while action serves either to bring sensory input in line with predictions or to actively obtain information that reduces uncertainty about task-relevant states (Parr et al., 2022; Priorelli et al., 2023; Russo et al., 2025). This view provides a principled account of why agents allocate gaze toward sources of uncertainty relevant to their current goals, as observed in the studies described above: gaze shifts can be understood as active, information-sampling decisions aimed at minimizing uncertainty about variables that matter for the task at hand, given the agent’s current predictive model of the task (Donnarumma et al., 2017; Itti & Baldi, 2009; Priorelli et al., 2025). In this perspective, the “uncertainty” and “priority” parameters of models such as the softmax barrier model are not merely descriptive quantities but can be grounded in the structure of the agent’s underlying generative model, opening the possibility that different people - using distinct generative models - could adopt qualitatively different gaze strategies for the same task.

Indeed, Decroix et al. (2017) showed that in alpine slalom skiing, expert skiers fixated on upcoming pole earlier than less skilled skiers. In additions, less skilled skiers spent more time looking at the surface between poles, suggesting that they sought information to better control their turn, whereas experts either did not need this information or relied on peripheral vision instead. More recently, we showed an even more profound difference in the visuomotor strategy between novice and skilled participants in a football task (Dehorne et al., 2025). Specifically, novices adopted a sequential approach of the task, dealing with actions one after another (i.e., looking at the ball while receiving the pass, then searching for passing opportunities, then passing the ball). Skilled participants, by contrast, embedded the visual search for the second action within the execution of the first action, enabling faster task completion (i.e., they looked at the ball and then directly searched for passing opportunities by looking at their teammate, before receiving and passing the ball). Within the “predictive brain” view introduced above, such differences would reflect distinct generative models of the task that would explain the differences in the sequence of sensorimotor decisions observed. Thus, conceiving sequential tasks as sequence of sensorimotor decisions involves deciding when to sample information for different subtasks, and these decisions appear to be optimized for performance outcomes (for example, to cope with the time pressure under which football players operate).

Experts and novices clearly differ in how they deploy their visual system, with distinct gaze strategies identified across numerous tasks, which supports experts anticipation skills (D. L. Mann et al., 2019; D. T. Y. Mann et al., 2007). These anticipation skills have been shown to rely on picking up more reliable cues at the right time, particularly when judging deceptive actions, on the ability to recognize patterns in evolving sequences of play, and on prioritizing some unfolding events or potential courses of action over others (Williams & Ford, 2008; Williams & Jackson, 2019; Yarrow et al., 2009). However, fundamental questions remain open to apprehend experts and novices differences in visuomotor behavior: how do eye movements and motor skills co-develop with practice (De Brouwer et al., 2021)? What factors set the timing of gaze shifts relative to movements (Hayhoe, 2017)? More fundamentally, what properties of the agent’s generative model give rise to qualitatively distinct visuomotor strategies across skill levels?

Recent computational work has begun to address this last question by formalizing the parameters of the generative model that distinguish expert from novice performers in anticipation tasks. Harris et al. (2022) showed that key findings from the sports anticipation literature can be accounted for within an active inference framework, and importantly argued that active inference can flexibly account for individual differences in anticipation through differences in prior beliefs and in sensory precision (i.e., the precision of the likelihood mapping between observations and hidden states). They also proposed that differences in preferred outcomes and observations could affect action selection. Relatedly, two studies demonstrated that fitting a Partially Observable Markov Decision Process (POMDP) model to expert and novice performances revealed differences in two parameters of their underlying generative model (Harris et al., 2023; He & Duan, 2025). Their results showed that expert performance was supported by higher sensory precision, and flexible prior beliefs about how the task unfolds. Interestingly, having flexible priors appeared advantageous for performance, which may seem at odds the idea that expertise is associated with prioritizing some courses of action over others, a pattern that would instead imply stronger, more confident priors (Williams & Jackson, 2019; Yarrow et al., 2009). This apparent tension may, however, reflect the nature of the task. Indeed, these two studies relied on a classic paradigm that consists in watching an occluded video and making a discrete choice to predict the outcome of the video, in which flexibility of prior beliefs is advantageous precisely because the task requires discriminating between a limited set of discrete outcomes (Harris et al., 2023; He & Duan, 2025). In sequential tasks requiring the coordination of multiple subtasks, such as the football task addressed here, the picture may be different: expert may hold stronger prior beliefs about how the task unfolds over time, enabling them to anticipate the informational demands of upcoming subtasks before they become immediately relevant, while remaining flexible at the level of individual action choices. It therefore remains an open question whether the parameters identified in discrete anticipation paradigm (i.e., high sensory precision and flexible structure of prior beliefs) can account for the qualitative shift from a sequential to a proactive visuomotor strategy observed in sequential tasks, where the challenge is not only to anticipate a single outcome, but to coordinate the timing of information sampling across multiple interdependent subtasks (Dehorne et al., 2025; Hayhoe & Ballard, 2014).

To address this open question, here we advance a “predictive brain” account of the football-task (Dehorne et al., 2025). Specifically, we implement computational models based on active inference, which provides a formalization of the “predictive brain” hypothesis, by casting both perception and action as processes that minimize free energy under an agent’s generative model (Friston, 2010; Parr et al., 2022). Free energy provides a tractable measure of surprise, reflecting the discrepancy between an agent’s predictions and its sensory observations. Under this framework, agents select policies, i.e., sequences of actions evaluated according to their expected free energy, thereby unifying uncertainty reduction (epistemic value) and reward maximization (pragmatic value) within a single objective function. In this sense, the active information-sampling decisions described above can be formally characterized as policies that minimize expected free energy under the agent’s generative model.

Our primary goal is providing a computational explanation of the differences between the strategies of novices and skilled participants, grounded in the active inference framework. For this, we design various active inference agents that address the football task using different parameterizations of generative models. Specifically, we manipulate three parameters of the agents’ generative models: the reward structure of the task, as novice and skilled participants are assumed to differ in how they value the different subtasks or their preferences about certain outcomes and observations; the prior beliefs about how the task unfolds over time; and sensory precision, defined by the mapping between hidden states and observations. Then, we perform a systematic model comparison, to assess which model parameters better reproduce the strategies of expert and novice participants. This model comparison will help us testing the relative roles of reward, prior beliefs and uncertainty in determining their observed differences in strategies.

## Methods

### Task and Simulation

In the real-world experiment that we address (Dehorne et al., 2025), the task consisted of receiving a football from a first teammate and then passing it to a second teammate. This task required participants to perceive how to receive the ball (e.g., on the right or left foot) and to determine the movement of the second teammate (e.g., running to the right or to the left) (Figure 1). For the purpose of modelling, this task was simplified by discretizing both the states and the actions into a finite set of possibilities. Thus, participants could be confronted with four possible conditions (i.e., two sides for receiving the ball and two possible running directions), for which they had no prior information. Importantly, participants could actively gather information about how to receive the ball (by looking at the first teammate and the ball trajectory) and to determine the movement of the second teammate (by looking at the second teammate), but they could not do both simultaneously. Therefore, the participants must decide whether to gather information (to reduce uncertainty before receiving and passing the ball) and which information to gather (about the first or second teammate, or both).

**Figure 1.**
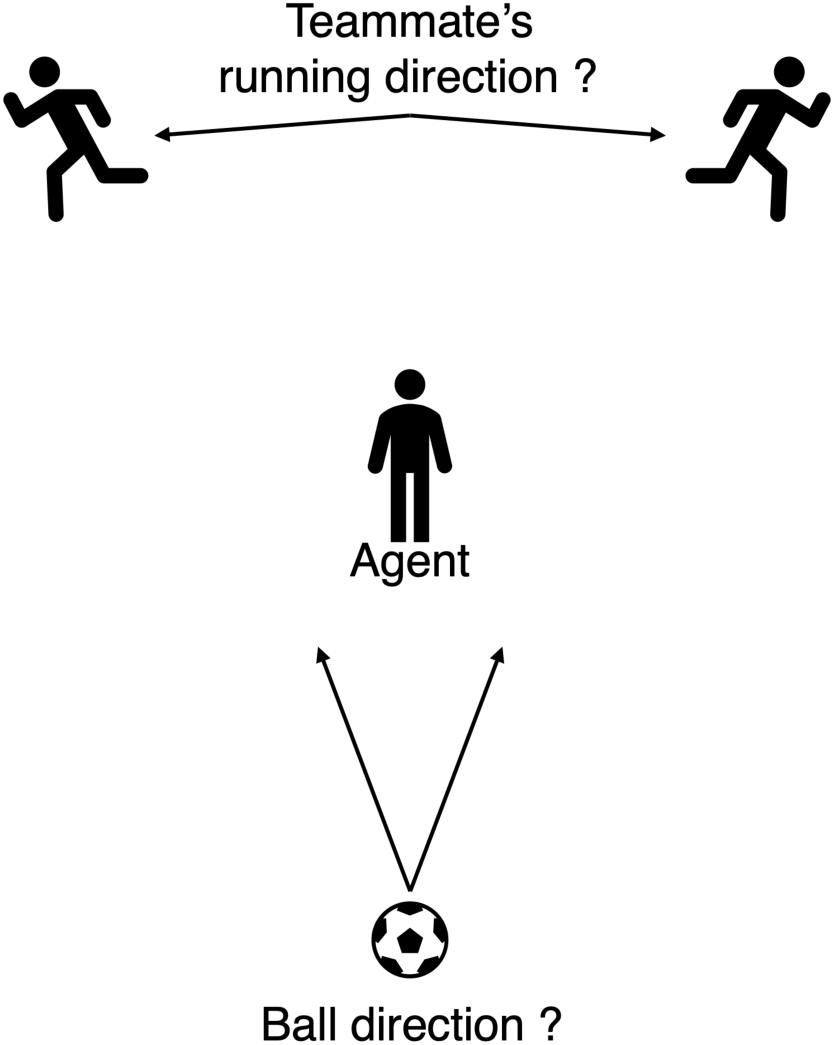
Representation of the experimental football task. The participant (in the central position) has to receive the ball from a first teammate and then pass it to a second teammate; for this, it can acquire information about the first or second teammate, or both.

The agent must act to identify both the ball reception side and the running direction of the teammate. These ambiguous decisions (actions) can be solved by looking at the ball or the teammate’s movement, respectively. Thus, these cues can’t be accessed simultaneously. In total, the agent can perform eight possible actions: looking at the ball going to the right or left, looking at the teammate running to the right or left, receiving the ball on the agent’s right foot, or left foot, passing the ball to the right or the left. A ninth action was included to the model, consisting in disengaging to the task.

To solve the task, we endowed the agent with five possible policies (i.e., sequences of actions). The first consists in looking at the ball twice (to increase confidence in the inferred state), then receiving the ball, looking at the running teammate, and finally passing the ball toward the running teammate. This policy corresponded to the strategy predominantly used by novice participants in the experiment (Dehorne et al., 2025). The second policy reflected the strategy used by expert players. It consisted in looking at the ball and looking at the running teammate, before receiving the ball and passing it to the running teammate. The remaining three policies were not empirically observed, but they were included in the model as plausible alternatives. The third and fourth policy consisted of looking toward only one of the two relevant areas (either the ball or the running teammate), followed by receiving the ball and directly passing it. The fifth was a purely pragmatic strategy: the agent directly received and passed the ball.

### Setup of the Generative Model

The active inference agent uses a probabilistic generative model, formalized as a POMDP model (Friston et al., 2016; Parr et al., 2022). The structure of the generative model for one trial is presented in Figure 3. Essentially, the generative model describes the probabilistic contingencies between (hidden) states (S), such as the position of the ball and of the teammate, actions or action sequences (policies *π*), such as looking at the ball or at the teammate position, and observations (O), over six timesteps (indexed by the subscripts). The first timestep corresponded to the beginning of the trial. During the next two timesteps, the agent could look at informational cues. Timestep four was the only time point at which the agent could receive the ball and obtain the associated reward. The final two timesteps allowed the agent either to pass the ball to the running teammate or to look at the teammate’s run. If the ball was passed at timestep five, the agent returned to the starting state at timestep six. These alternative action sequences correspond to five action sequences (or policies) which are formally encoded in the V^1^-matrix over these six timesteps (Figure 3). They correspond to the five policies described above (i.e., the novice, expert, looking only at the ball, looking only at the teammate and pragmatic strategies). Their prior probabilities were defined in the V^2^-matrix, with all policies assigned equal prior probability at the beginning of each trial. In our simulation, the agent could not update their prior beliefs from trial to trial.

At timestep *t*, agent has observation *o_t_*, while performing an action *a_t_* generated from policy *π* and state *s_t_*. The generative model presents as:

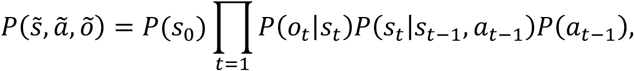

To relate this equation to the POMDP structure in Figure 2, *P*(*s*_0_) represents the priors over initial states D, *P*(*o_t_*|*s_t_*) the likelihood mapping between states and observations A, and *P*(*s_t_*|*s_t_*_–1_, *a_t_*_–1_) the beliefs about state transitions B.

**Figure 2.**
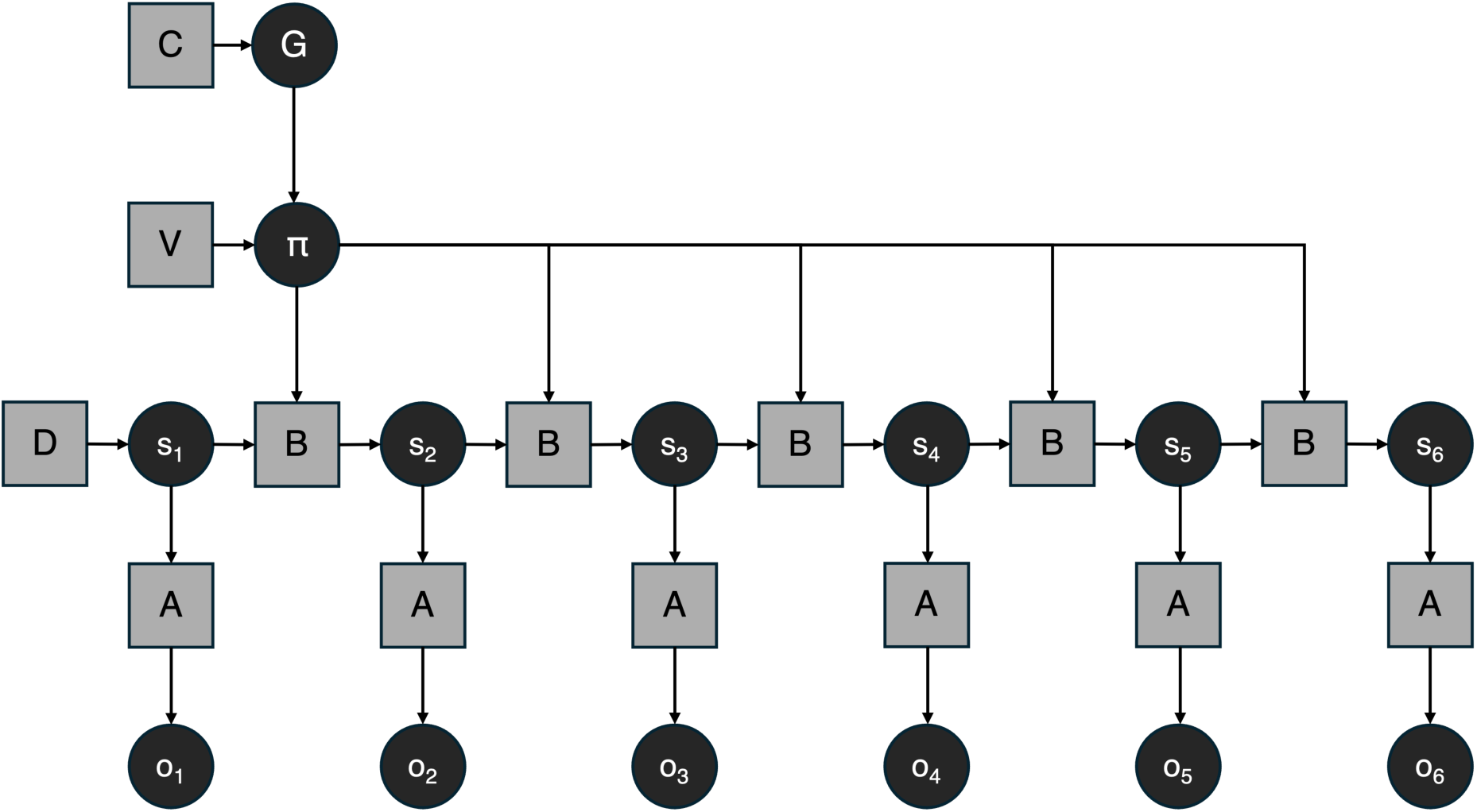
Structure of the partially-observable Markov decision process model (POMDP) used for the football task. The circular nodes represent random variables, whereas squares represent parameters of the generative model. Hidden state (s) generate observations (o) at each of the six timesteps of the trial. A represents the likelihood mapping between states and observations. B specifies the beliefs about state transitions, which encodes action-dependent state dynamics. D contains the priors about initial states. Policies (π) are sequence of actions, defined by V, that are selected based on their expected free energy (G). C contains the prior preferences over observations that influence policy selection through G.

Policy selection will depend on a probability distribution based on expected free energy *G*. The lower this value, the higher the probability of being selected. Expected free energy represents how likely the policy *π* will generate the agent’s preferred observations at a given timestep *t*:

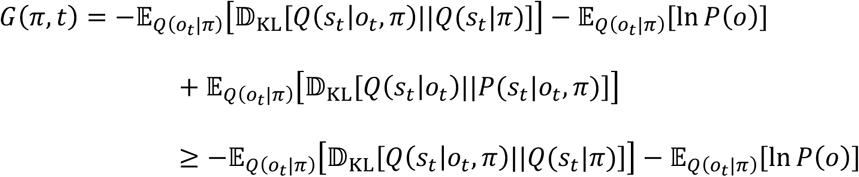

This shows that expected free energy can be decomposed into an epistemic value, representing the information gain between the prior *Q*(*s_t_*|*π*) and posterior *Q*(*s_t_*|*o_t_*, *π*) using a Kullback-Leibler divergence measure, and a pragmatic value representing the expected log probability of observing the preferred observation *P*(*o*) under the selected policy. In the POMDP structure (Figure 2), the preferences over observations are set in C. Therefore, the approximate posterior of the policy *Q*(*π*) is proportional to the expected free energy in a softmax function *σ*:

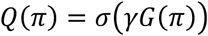

The expected free energy *G* represents the agent’s variational free energy expected after pursuing a policy *π,* and is therefore a measure of free energy in the future. Free energy is also evaluated based on past/present observations with the variational free energy *F*. The agents minimize the variational free energy *F* through perception (by updating beliefs about current states) and by selecting the policy with the lowest *G*. The variational free energy evaluates the discrepancy (or Bayesian surprise) between the joint distribution *P*(*s*~, õ,ã) and the approximate posterior distribution (here represented by *Q*(*s*~, õ,ã)):

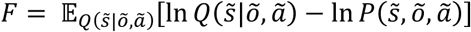

Each parameter of the generative model (squares in Figure 2) was specified as follows for the present task. The D vectors present the probability of starting in a given state and the probability of being in a given context. In our task, agents were always starting in the starting position, for which the first state was associated. Thus, the first D vector take the following form:

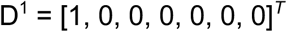

We also set the second D vector so that the four contexts had the same probability of occurrence and could not be updated from trial to trial:

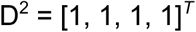

A-matrices encode the likelihood mapping between states and observations, that is, the probability of observing an outcome given a hidden state. In our model, this mapping included both exteroceptive (A^1^-matrices) and interoceptive (A^2^-matrices) modalities (Figure 3). Depending on the context of the trial, the hidden states related to attending to the ball or to the teammate’s run generated distinct observations. These observations could then inform action selection, enabling the agent to correctly receive the ball and pass it in the appropriate direction. We hypothesized that reducing the precision of the likelihood mapping would capture novice behavior. In this framework, lower likelihood precision corresponds to lower sensory precision, that is, making observations that are less reliable for inferring the underlying hidden state. For example, this would correspond to a novice player visually tracking the ball, but using ambiguous or noisy cues about its trajectory. To test this hypothesis, we manipulated the precision of the likelihood mapping by adding noise in the A-matrices. We expected that decreasing sensory precision would bias policy selection toward the novice strategy.

**Figure 3.**
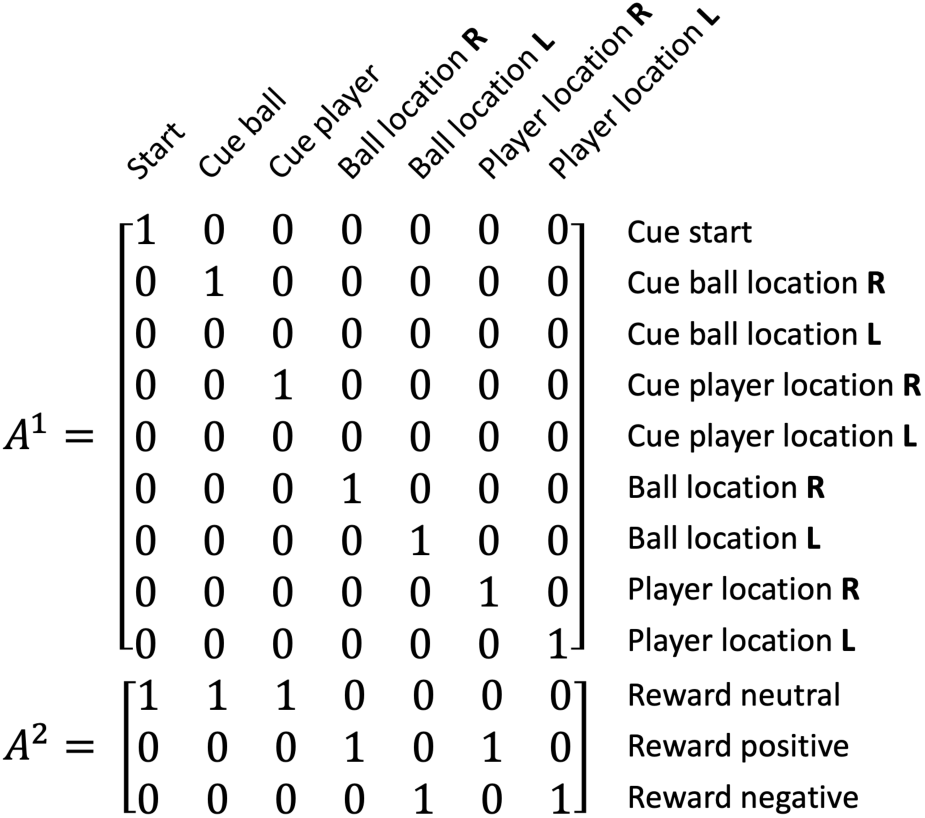
Example of A-matrices for one of the 4 contexts, here with the ball coming on the right and the running player going to the right. The first matrix presents the probability mapping of states with exteroceptive cues, and the second with interoceptive cues.

B^1^-matrices encodes the state transition probabilities, that is, the probability of transitioning to a subsequent hidden state given the current state and the selected action (Figure 4). These matrices therefore reflect the agent’s beliefs about how actions influence state dynamics. By introducing noise into B^1^-matrices, we increased uncertainty about action-dependent state transitions, that is, uncertainty regarding the consequences of the agents’ actions. We hypothesized that a lower transition precision would capture key aspect of novice performance. In behavioral terms, novice participants lower technical skill may render action outcomes less predictable. For example, the direction of their pass could occasionally deviate from the intended one. To test this hypothesis, we manipulated the precision of the transition probabilities and predicted that increasing uncertainty about action outcomes would bias policy selection toward the novice strategy.

**Figure 4.**
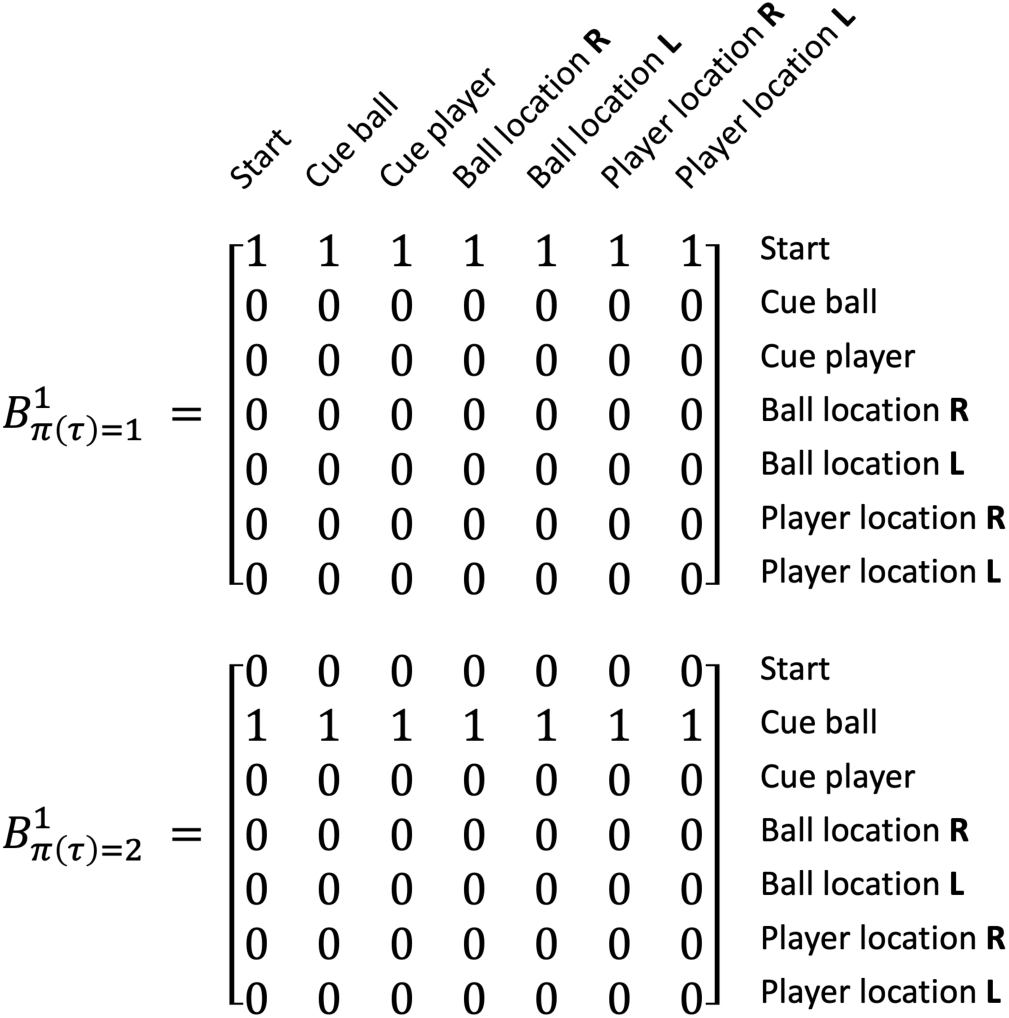
Example of B-matrices for two of the seven possible actions. The matrices present the transition probabilities for transition between the possible states. The first matrix corresponds to moving to the starting location. The second matrix corresponds to obtaining cue about the ball location. In both cases, agents are certain to be in a particular state on the next timestep due to their action.

A second B-matrix encodes the probability of transitions between contexts from one timestep to the next within a trial (Figure 5). In our task, context remained constant over time, such that this matrix was an identity matrix.

**Figure 5.**
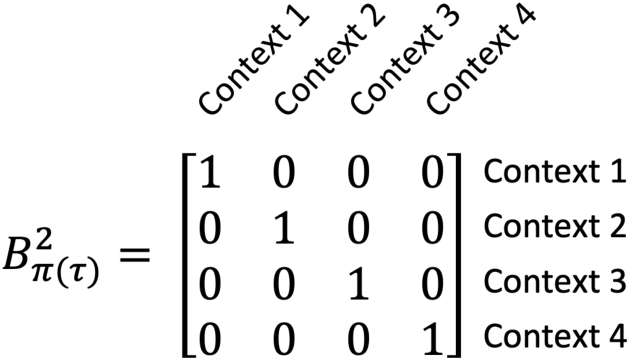
Presentation of the B-matrix about context transition. It shows that context stays constant over time.

The C-matrices encode the prior preferences over observations across the six timesteps of each trial (Figure 6). These preferences specify the relative value assigned to different outcomes and shape policy selection through the minimization of expected free energy, primarily via its pragmatic (reward-seeking) component (Parr et al., 2022). In the first C matrix (C^1^ in Figure 6), we specified a strong preference for receiving the ball at timestep four. This constraint was introduced to simulate the requirement that the agent commit to a decision regarding the ball reception at that specific time point, and prior to pass the ball. The goal was to approximate the temporal structure of the real-world task. Then, the agents had two possible timesteps (five and six) to pass the ball. This temporal window enabled two distinct strategies: acting immediately (as generally observed in skilled players) or looking at the teammate’s position before passing the ball (as more generally observed in the novices). We hypothesized that skilled performers would exhibit a stronger preference for rapid action, reflecting their capacity to operate efficiently under the spatiotemporal constraints imposed by opponents in real-game situations. Within the Active Inference framework, increasing the reward value associated with passing at time step five raises the expected utility of immediate action. In this sense, higher positive reward values can be interpreted as promoting a form of risk-seeking behavior, as the agent becomes more inclined to commit to an early pass without fully resolving uncertainty about the teammate’s position. Conversely, lower reward values reduce the incentive for immediate action and relatively increase the contribution of epistemic value, thereby favoring information-seeking strategies. To test this hypothesis, we manipulated the value of the reward at time step five in the second preference matrix (C^2^ in Figure 6), expecting that it would shift the balance between rapid, skilled-like performance, and sequential, information-seeking (novice-like) behavior.

**Figure 6.**
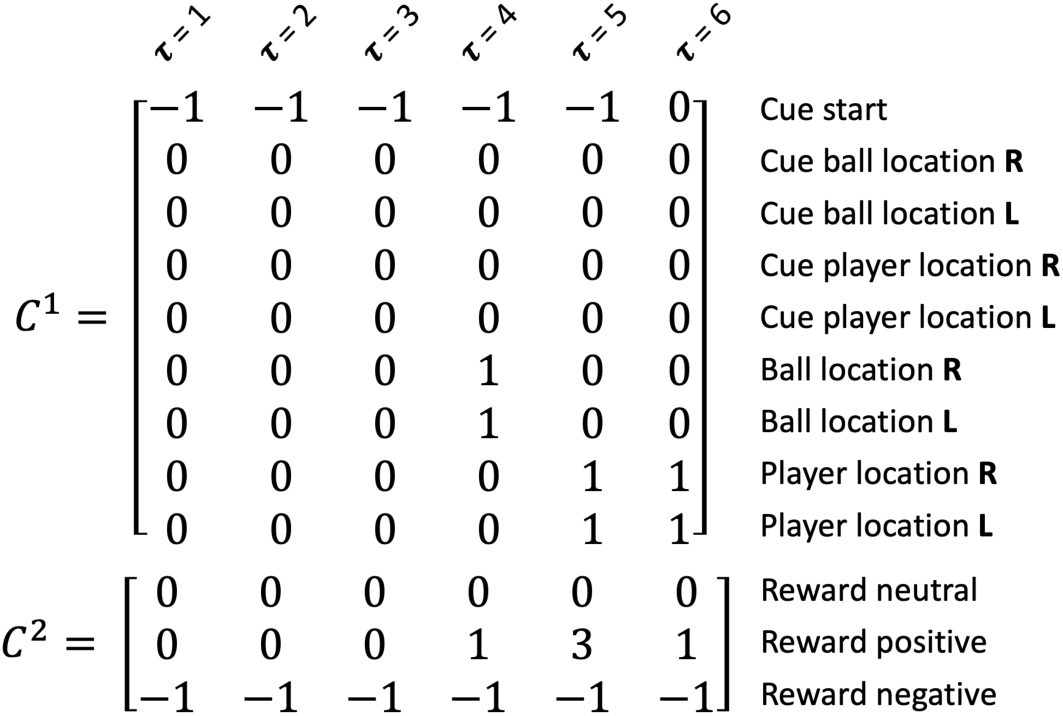
Presentation of the two C-matrices. The C^1^ matrix presents the prior preferences over outcomes. Agent does not like to be at the starting position at timesteps 1-5. The agent likes to be at the ball locations at timestep 4, and at the player locations at timesteps 5-6. The C^2^ matrix presents the prior preferences for neutral, attractive and aversive stimuli. The positive stimulus becomes attractive at timesteps 4, 5 and 6. But, its higher value on timestep 5 compared to 6 should promote passing the ball at timestep 5.

### Procedure of the Simulations

We used the generative model described above to simulate policy selection in the football task. For each parameter configuration, simulations were run with 50 virtual agents, each performing 40 trials (10 trials in each of the four contexts, presented in randomized order). All simulations were implemented in MATLAB R2025b using the SPM12 toolbox.

As a complete exploration of all the parameter space would be very time-consuming, we decided to change one parameter at a time, starting with the one that was expected to produce the greatest change. In a first set of simulations, therefore, we manipulated the prior preferences over outcomes (C^2^-matrix). Specifically, we progressively increased the reward value at timestep five. The reward value was varied from 1 (indicating no preference between passing the ball at timestep five or six) to 6 (strong preference for passing the ball immediately after receiving it). Based on the results of this set of simulations, three representative levels of prior preferences were selected for subsequent simulations related to (i) beliefs about action consequences and, (ii) sensory precision.

Transition precision was progressively reduced by introducing noise into the B^1^-matrices. We began with maximal precision, corresponding to the deterministic transition structure shown in Figure 4. We then reduced the probability of the most likely state transition by .02 across four steps (i.e., probabilities of 1 were replaced by .98, .96, .94, and .92, respectively). The remaining probability was distributed across alternative transitions such that each column of the transition matrices continued to sum to one.

Sensory precision over exteroceptive signal was progressively lowered by introducing noise in the A^1^-matrices. As with the transition matrices, we began with a noise-free likelihood mapping (Figure 3) and applied the same procedure as described for the B^1^-matrices, reducing the probability of the most likely observation and redistributing probability across alternative outcomes.

For each parameter configuration, we recorded (i) the sequence of action performed by the virtual agent (i.e., the selected policy on each trial), (ii) decision accuracy (i.e., whether the agent successfully received the ball and passed it to the teammate) and (iii) the total reward accumulated over the 40 trials, expressed relative to the maximum achievable reward under the specified prior preferences.

## Results

### Manipulation of Prior Preferences over Observations

We progressively increased, in the C^2^ matrix, the reward value for passing the ball at timestep five. With respect to policy selection, we observed a clear threshold at a reward value of 3 (Figure 7A). For reward values of 1 and 2, virtual agents exclusively selected the novice strategy. In contrast, for reward values ranging from 3 to 6, agents exclusively selected the expert strategy, characterized by rapid passing following ball reception.

**Figure 7.**
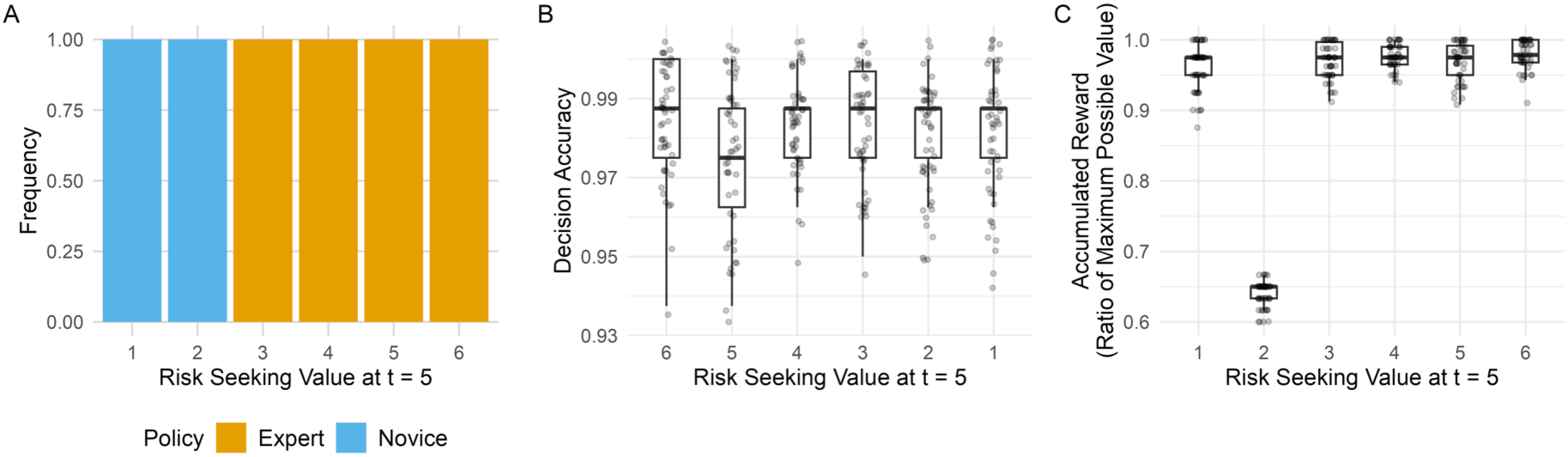
Policy selection (A), decision accuracy (B) and reward obtained (C) under different values of positive reward at timestep five.

The shift in strategy did not significantly affect decision accuracy across conditions (Figure 7B), *F*(5,294) = 1.64, *p* = .149, *η^2^* = .03. Overall, agents successfully received the ball and passed it in the correct direction in the vast majority of trials (*M* = 98.1%, *SD* = 1.5%).

However, when the reward value at time step five was set to 2, selection of the novice strategy was suboptimal in terms of total reward obtained relative to the maximum achievable reward (*M* = 63.9%, *SD* = 1,8%, Figure 7C). This performance was significantly lower than in all other conditions, *F*(5,294) = 1617, *p* < .001, *η^2^* = .97 (all post hoc comparisons between the reward value of 2 and other conditions: *ps* < .001 and *ds* > 13). Two additional, comparatively small post hoc differences were observed between the reward value of 1 and reward values of 4 (*p* = .017, *d* = 0.66) and 6 (*p* = .002, *d* = 0.79). Detailed results of the post hoc tests are provided in the Supplementary Material.

Based on these results, we selected three representative levels of prior preferences for subsequent simulations: (i) a constant reward value across timesteps, (ii) a medium reward value at timestep five (value of 3), promoting rapid performance and consistent selection of the expert strategy, and (iii) a high reward value at timestep five (value of 5), strongly favoring immediate passing and the expert-like behavior.

### Manipulation of Beliefs about Transition Dynamics

For the three selected levels of positive reward at timestep five, we progressively reduced transition precision by introducing noise into the B^1^-matrices. When reward values were held constant across timesteps, the novice strategy remained the dominant policy, even at the highest levels of transition noise (Figure 8A). However, increased noise induced occasional stochastic selection of alternative policies across trials, suggesting reduced confidence in action-dependent state transitions.

**Figure 8.**
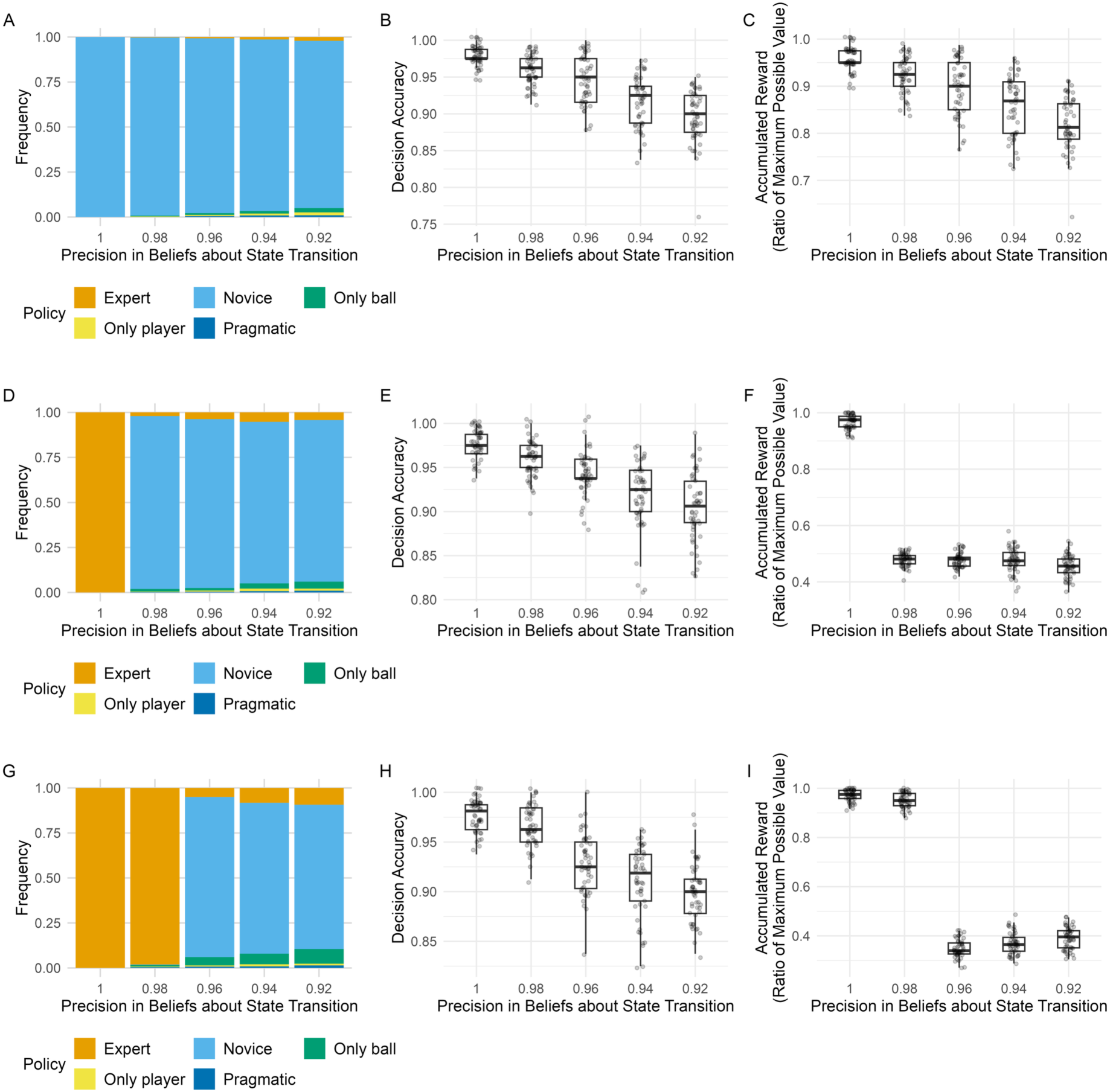
Policy selection, decision accuracy and reward obtained under different level of noise in beliefs bout action consequences. Panels A, B and C present results under constant value of positive reward. Panels D, E and F show results under medium reward value at timestep five. Panels G, H and I show results under high reward value at timestep five.

Under medium and high reward values at time step five, reducing transition precision induced a clear shift in policy selection (Figure 8D and 8G, respectively). Specifically, agents initially selected the expert strategy under high precision, but progressively reverted to the novice strategy as transition noise increased. Notably, this transition occurred at higher noise levels under the high-reward condition compared to the medium-reward condition (Transition precision of .96 and .98, respectively), indicating that stronger prior preferences for rapid passing partially compensated for reduced confidence in action outcomes.

Increasing transition noise significantly impaired decision accuracy, irrespective of the level of reward manipulation (Figure 8B, 8E and 8H). Accuracy declined monotonically as noise increased, dropping from approximately 98% in the absence of noise to 90% at the lowest level of precision (.92). Detailed statistics of the ANOVAs are provided in Table 1, and results of the post hoc tests are provided in the Supplementary Material.

**Table 1.** One-way analyses of variance in decision accuracy and proportion of reward collected under the three levels of prior preferences for passing at timestep 5.

| Measure and Level of Prior Preference at $\tau = 5$ | $F(4,245)$ | $p$ | $\eta^2$ |
| --- | --- | --- | --- |
| Decision Accuracy |  |  |  |
| Constant reward | 68.6 | < .001 | .53 |
| Medium reward | 50.3 | < .001 | .45 |
| High reward | 81.1 | < .001 | .57 |
| Proportion of Reward Collected |  |  |  |
| Constant reward | 58.7 | < .001 | .49 |
| Medium reward | 2294 | < .001 | .97 |
| High reward | 3909 | < .001 | .99 |

With constant reward values, the proportion of reward collected decreased as transition noise increased (Figure 8C), consistent with the observed decline in decision accuracy. In contrast, under medium and high reward conditions (i.e., when risk-seeking was promoted), reward collection exhibited a sharp decline when agents shifted away from the expert strategy toward the novice strategy (Figure 8F and 8I, respectively). This reduction reflects the temporal structure of outcome preferences: under these conditions, passing at timestep six remained rewarding but yielded a lower value than passing at timestep five. Consequently, deviations from rapid action were disproportionately penalized in high reward contexts. Results also revealed that the proportion of reward collected significantly increased from transition precision value of .96 to .92. This appeared also related to increased proportion of selection of policy related to rapid action.

### Manipulation of Sensory Precision

When introducing noise into the A^1^-matrices across the three levels of risk-seeking at timestep five, a distinct pattern of policy selection emerged compared to the manipulation of transition precision. Under constant value of reward, the novice strategy remained the dominant policy regardless of the level of sensory precision (Figure 9A). However, decreasing sensory precision up to .94 led to a progressive decline in decision accuracy (Figure 9B) and in the proportion of reward collected (Figure 9C), consistent with reduced reliability of sensory evidence for inferring hidden states. Detailed statistics of the ANOVAs are provided in Table 2, and results of the post hoc tests are provided in the Supplementary Material.

**Figure 9.**
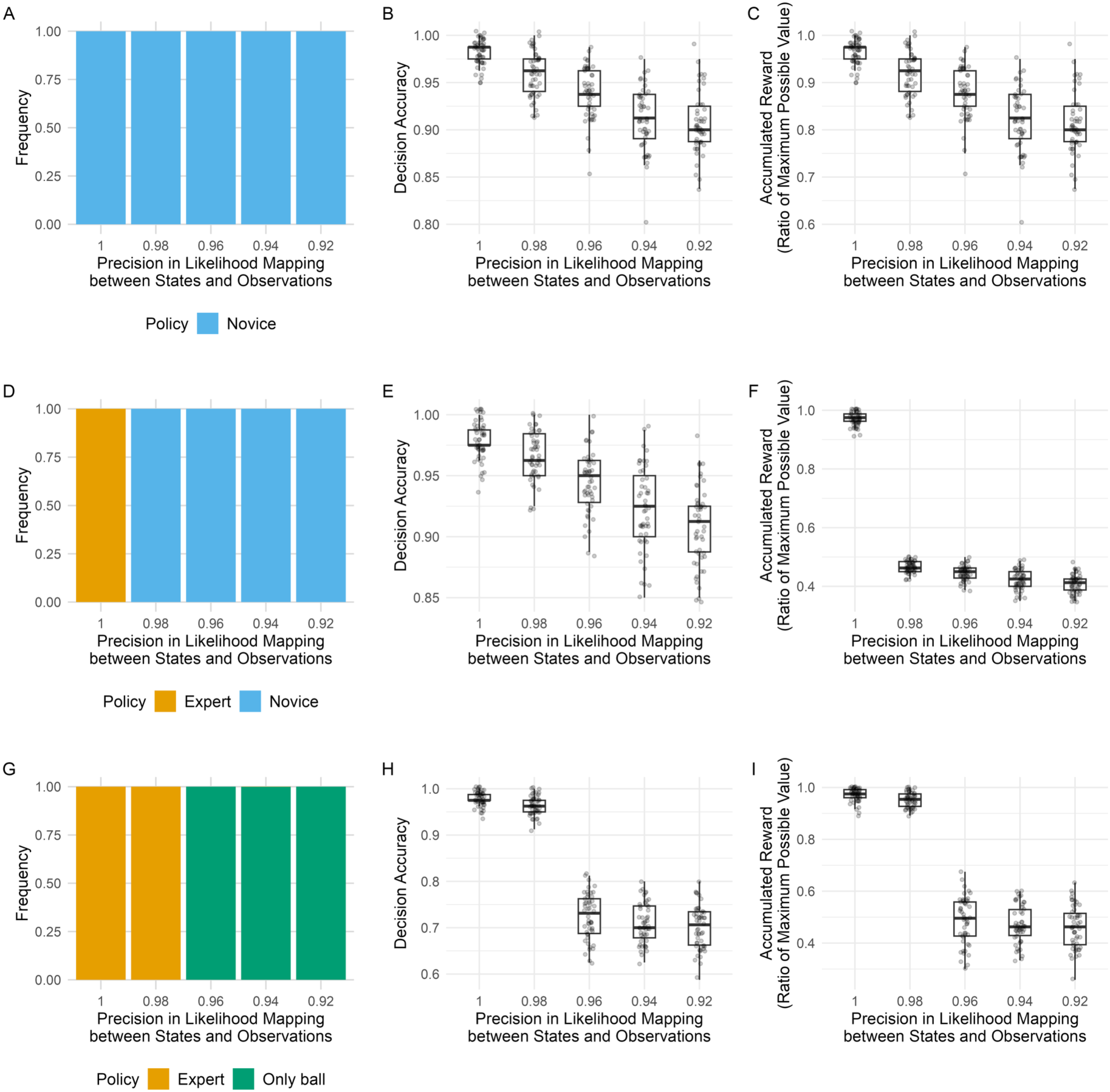
Policy selection, decision accuracy and reward obtained under different level of sensory precision. Panels A, B and C present results under constant value of positive reward. Panels D, E and F show results under medium reward value at timestep five. Panels G, H and I show results under high reward value at timestep five.

**Table 2.** One-way analyses of variance in decision accuracy and proportion of reward collected under the three levels of prior preferences for passing at timestep 5.

| Measure and Level of Prior Preference at $\tau = 5$ | $F(4,245)$ | $p$ | $\eta^2$ |
| --- | --- | --- | --- |
| Decision Accuracy |  |  |  |
| Constant reward | 71.3 | < .001 | .54 |
| Medium reward | 62.7 | < .001 | .51 |
| High reward | 726 | < .001 | .92 |
| Proportion of Reward Collected |  |  |  |
| Constant reward | 71.3 | < .001 | .54 |
| Medium reward | 3875 | < .001 | .98 |
| High reward | 830 | < .001 | .93 |

Under the medium reward condition, even the lowest level of perceptual noise tested was sufficient to induce a shift from the expert strategy to the novice strategy (Figure 10D). This transition in policy selection was accompanied by a marked decrease in the proportion of reward collected (Figure 9F). After this transition, the proportion of reward collected continued to decrease as the sensory precision decreased, but with a lower magnitude. Decision accuracy also progressively declined with sensory precision (Figure 9E).

Finally, under the high reward condition, increasing perceptual noise also led to a transition away from the expert strategy (Figure 9G). However, in this case, the alternative policy selected was not the fully novice strategy. Instead, agents adopted an intermediate strategy: looking only at the ball trajectory prior to receiving the ball, and then passing immediately at timestep five without looking at the teammate’s running direction. This shift suggests a reconfiguration of the exploration–exploitation balance under high preference for immediate passing. As perceptual precision decreased, agents appeared to preserve rapid action (to secure the higher reward at time step five) while reducing looking at the teammate’s movement. This change in strategy was associated with a significant drop in both decision accuracy (Figure 9H) and proportion of reward collected (Figure 9I), particularly at the noise level corresponding to the policy transition.

## Discussion

The present study aimed to model the visuomotor strategies observed in a sequential football task using an active inference framework. Specifically, the model was designed to reproduce the generative models that would lead to the selection of two qualitatively distinct policies: a novice strategy, in which visual information about each subtask is obtained only when immediately required, and a skilled strategy, in which information about upcoming subtasks is obtained proactively during the execution of the current one. Simulations successfully lead to the selection of one of the two policies, and identified three key parameters driving the preference for one policy over the other: the reward structure of the task, the precision in state transitions, and the level of sensory precision. These settings of the generative model also affected the task performance in terms of accuracy. More precisely, simulations showed that the selection of the expert policy relied on strong prior beliefs about state transition, high sensory precision and high valuation of fast passing after ball reception. These settings were also associated with high decision accuracy. Conversely, the selection of the novice policy was selected in particular when rapid performance was not highly valued, or under lower precision in state transitions, or lower level of sensory precision, which also decreased decision accuracy. Taken together, these results suggest that the policy selection leading to performing the novice or the skilled visuomotor strategy in this sequential task reflects differences in the underlying generative model, particularly in how agents value different aspects of the task.

The temporal structure of rewards had a key role in our simulations. Given the spatiotemporal pressure experienced by football players in performance context, we expected that skilled players would assign a higher expected utility to fast passing after ball reception. This was effectively affecting policy selection, with skilled strategies being more consistently selected when fast response was more rewarded. This is consistent with previous models showing that visuomotor behavior was shaped by assigned task priority (Johnson et al., 2014; Rothkopf et al., 2016). However, our results add that the temporal reward structure of the task can shape the sequence of sensorimotor decisions in a proactive manner, such that epistemic actions like looking toward a particular location can be anticipated before they are immediately necessary, in order to secure higher reward at a later stage of the task. Thus, the sequence of pragmatic and epistemic actions appears to be shaped by a temporal structure of reward that has to be acquired during practice. From a theoretical view, this potential shift in the structure of rewards echoes a learning process from ecological psychology called education of intention (Arzamarski et al., 2010; Jacobs & Michaels, 2007). This process propose that learners change their goals with practice so that they converge with the task-goals. Intentions are closely related to perception and action, as they closely affect their utility from the agent perspective thus guiding their exploratory behavior. Being able to model the temporal structure of reward based on observed behavioral data could help capture this shift in intentions with practice to further understanding of perceptual-motor skill acquisition, which generally assumes that agent’s intentions directly match task structure and instructions.

Our simulations showed that acquiring this specific temporal reward structure is not sufficient for the selection of the skilled strategy. It also requires high confidence in state transitions and high sensory precision, consistent with the notion that expertise refines the generative model underlying action selection. High confidence in state transitions implies that the agent can reliably anticipate the informational demands of upcoming subtasks before they become immediately relevant, which appears to be key aspect of the proactive strategy observed in skilled participants (Dehorne et al., 2025). This is consistent with the literature showing that with practice, knowledge about environmental event statistics can be acquired to optimize the timing of gaze shifts (Hoppe & Rothkopf, 2016), or blinks (Hoppe et al., 2018). This proactive use of the generative model also resonates with the affordance competition hypothesis (Cisek, 2007; Cisek & Kalaska, 2010; Pezzulo & Cisek, 2016), according to which action selection is biased not only by the immediate outcomes of currently available affordances but also by predictions of what new affordances they will make available, allowing agents to purposively create future opportunities for action. The proactive strategy observed in skilled players can be understood in these terms: by obtaining visual information about the second subtask during the first, skilled players actively create the conditions for a subsequent affordance to become available earlier, rather than merely reacting to it once the first subtask is complete. This interpretation is supported by recent evidence from a sequential embodied decision task, in which participants who assigned greater weight to future outcomes achieved better performance and made fewer errors than those who relied primarily on immediate affordances (Nuzzi et al., 2026), and in which deliberation continued throughout movement execution rather than being completed beforehand, suggesting that beliefs about state transitions and sensory precision are continuously exploited and refined as the task unfolds, consistent with the active, online use of the generative model proposed here.

Beyond reproducing the selection of the observed behavioral strategies, the active inference framework offers a principled account of how perception and action are jointly organized in sequential tasks. Previous models of visuomotor behavior in multitask settings relied on reward-based or priority-based accounts of gaze allocation (Johnson et al., 2014; Rothkopf et al., 2016), successfully capturing how agents allocate gaze across competing sources of information. In parallel, POMDP models applied to anticipation skills in occlusion paradigms suggested that expertise was characterized by flexible priors to adapt their decision to informational cues (Harris et al., 2023; He & Duan, 2025). However, these models did not address how agents decide when to shift gaze relative to movement performance in a sequential context. By integrating both epistemic and pragmatic actions in the process of policy selection through minimization of free energy under a generative model, active inference provides a unified account of these decisions: the timing of epistemic actions relative to pragmatic ones emerges naturally from the interaction between the agent’s beliefs about task structure, its preferences over observations, and its uncertainty about the current state. In this sense, active inference reframes the problem of sequential sensorimotor decisions as one of inference under a generative model, where the timing of epistemic actions is not prescribed but emerges from the structure of the agent’s beliefs and preferences.

The current study has several limitations. First, the task was simplified for modelling reasons, in particular, behaviors and conditions were discretized as required POMDP. Therefore, it doesn’t take into account the larger panel of behavior that this kind of task offers and the more complex constraints that apply on policy selection. Second, an alternative explanation of policy selection can be related to habit formation, with skilled strategy possibly being reinforced during practice. This would be possible in the simplified version of the task presented here, but given the current pattern of the results, we would expect that in a more complex situation, experts would scale the epistemic actions to expected time available in the unfolding context, providing them with more behavioral flexibility to adapt to different demands. For example, receiving a ball from further away may provide enough time to look toward more than one available teammate demanding the ball, but this requires an effective set of beliefs about how and when the ball will be received based on information obtained from looking at the departure of the ball. These expectations might be testable experimentally through the manipulation of the passing distance, for example.

