## Supplementary Material for "Expert and Novice Visuomotor Strategies in a Sequential Football Task Reflect Distinct Generative Models"

**Table S1**

Results of the post hoc tests for the proportion of reward collected when manipulating prior preferences over outcomes.

| Reward Value at $\tau = 5$ | | | | | | | | |
|---|---|---|---|---|---|---|---|---|
| condition | | condition | Mean difference | Standard Error | dof | t | $p_{bonferroni}$ | Cohen's d |
| 1 | - | 2 | 0.32200 | 0.00478 | 294 | 67.352 | <.001 | 13.4705 |
| | - | 3 | -0.00950 | 0.00478 | 294 | -1.987 | .718 | -0.3974 |
| | - | 4 | -0.01570 | 0.00478 | 294 | -3.284 | .017 | -0.6568 |
| | - | 5 | -0.00883 | 0.00478 | 294 | -1.848 | .985 | -0.3695 |
| | - | 6 | -0.01886 | 0.00478 | 294 | -3.944 | .002 | -0.7889 |
| 2 | - | 3 | -0.33150 | 0.00478 | 294 | -69.340 | <.001 | -13.8679 |
| | - | 4 | -0.33770 | 0.00478 | 294 | -70.636 | <.001 | -14.1273 |
| | - | 5 | -0.33083 | 0.00478 | 294 | -69.200 | <.001 | -13.8400 |
| | - | 6 | -0.34086 | 0.00478 | 294 | -71.297 | <.001 | -14.2594 |
| 3 | - | 4 | -0.00620 | 0.00478 | 294 | -1.297 | 1.000 | -0.2594 |
| | - | 5 | 6.67e-4 | 0.00478 | 294 | 0.139 | 1.000 | 0.0279 |
| | - | 6 | -0.00936 | 0.00478 | 294 | -1.957 | .769 | -0.3914 |
| 4 | - | 5 | 0.00687 | 0.00478 | 294 | 1.436 | 1.000 | 0.2873 |
| | - | 6 | -0.00316 | 0.00478 | 294 | -0.660 | 1.000 | -0.1321 |
| 5 | - | 6 | -0.01002 | 0.00478 | 294 | -2.097 | .553 | -0.4193 |

Note. Comparisons based on estimated marginal means.

**Table S2**

Results of the post hoc tests for the decision accuracy when manipulating transition precision (under constant reward value).

| Transition Precision | | | | | | | | |
|---|---|---|---|---|---|---|---|---|
| condition | | condition | Mean difference | Standard Error | dof | t | $p_{bonferroni}$ | Cohen's d |
| 0.92 | - | 0.94 | -0.0225 | 0.00565 | 245 | -3.98 | <.001 | -0.797 |
| | - | 0.96 | -0.0487 | 0.00565 | 245 | -8.63 | <.001 | -1.726 |
| | - | 0.98 | -0.0635 | 0.00565 | 245 | -11.24 | <.001 | -2.248 |
| | - | 1.00 | -0.0837 | 0.00565 | 245 | -14.82 | <.001 | -2.965 |
| 0.94 | - | 0.96 | -0.0263 | 0.00565 | 245 | -4.65 | <.001 | -0.929 |
| | - | 0.98 | -0.0410 | 0.00565 | 245 | -7.26 | <.001 | -1.451 |
| | - | 1.00 | -0.0613 | 0.00565 | 245 | -10.84 | <.001 | -2.168 |
| 0.96 | - | 0.98 | -0.0147 | 0.00565 | 245 | -2.61 | .096 | -0.522 |
| | - | 1.00 | -0.0350 | 0.00565 | 245 | -6.20 | <.001 | -1.239 |
| 0.98 | - | 1.00 | -0.0203 | 0.00565 | 245 | -3.58 | .004 | -0.717 |

Note. Comparisons based on estimated marginal means.

**Table S3**

Results of the post hoc tests for the proportion of reward collected when manipulating transition precision (under constant reward value).

| Transition Precision | | | | | | | | |
|---|---|---|---|---|---|---|---|---|
| condition | | condition | Mean difference | Standard Error | dof | t | $p_{bonferroni}$ | Cohen's d |
| 0.92 | - | 0.94 | -0.0388 | 0.0101 | 245 | -3.84 | .002 | -0.768 |
| | - | 0.96 | -0.0783 | 0.0101 | 245 | -7.75 | <.001 | -1.550 |
| | - | 0.98 | -0.1050 | 0.0101 | 245 | -10.40 | <.001 | -2.080 |
| | - | 1.00 | -0.1395 | 0.0101 | 245 | -13.82 | <.001 | -2.763 |
| 0.94 | - | 0.96 | -0.0395 | 0.0101 | 245 | -3.91 | .001 | -0.782 |
| | - | 0.98 | -0.0662 | 0.0101 | 245 | -6.56 | <.001 | -1.312 |
| | - | 1.00 | -0.1007 | 0.0101 | 245 | -9.98 | <.001 | -1.996 |
| 0.96 | - | 0.98 | -0.0268 | 0.0101 | 245 | -2.65 | .086 | -0.530 |
| | - | 1.00 | -0.0613 | 0.0101 | 245 | -6.07 | <.001 | -1.213 |
| 0.98 | - | 1.00 | -0.0345 | 0.0101 | 245 | -3.42 | .007 | -0.683 |

Note. Comparisons based on estimated marginal means.

**Table S4**

Results of the post hoc tests for the decision accuracy when manipulating transition precision (under medium reward value).

| Transition Precision | | | | | | | | |
|---|---|---|---|---|---|---|---|---|
| condition | | condition | Mean difference | Standard Error | dof | t | $p_{bonferroni}$ | Cohen's d |
| 0.92 | - | 0.94 | -0.0137 | 0.00596 | 245 | -2.31 | .219 | -0.461 |
| | - | 0.96 | -0.0382 | 0.00596 | 245 | -6.42 | <.001 | -1.283 |
| | - | 0.98 | -0.0547 | 0.00596 | 245 | -9.18 | <.001 | -1.837 |
| | - | 1.00 | -0.0737 | 0.00596 | 245 | -12.37 | <.001 | -2.474 |
| 0.94 | - | 0.96 | -0.0245 | 0.00596 | 245 | -4.11 | <.001 | -0.822 |
| | - | 0.98 | -0.0410 | 0.00596 | 245 | -6.88 | <.001 | -1.375 |
| | - | 1.00 | -0.0600 | 0.00596 | 245 | -10.06 | <.001 | -2.013 |
| 0.96 | - | 0.98 | -0.0165 | 0.00596 | 245 | -2.77 | .061 | -0.554 |
| | - | 1.00 | -0.0355 | 0.00596 | 245 | -5.95 | <.001 | -1.191 |
| 0.98 | - | 1.00 | -0.0190 | 0.00596 | 245 | -3.19 | .016 | -0.637 |

Note. Comparisons based on estimated marginal means.

**Table S5**

Results of the post hoc tests for the proportion of reward collected when manipulating transition precision (under medium reward value).

| Transition Precision | | | | | | | | |
|---|---|---|---|---|---|---|---|---|
| condition | | condition | Mean difference | Standard Error | dof | t | $p_{bonferroni}$ | Cohen's d |
| 0.92 | - | 0.94 | -0.02000 | 0.00652 | 245 | -3.067 | .024 | -0.6134 |
| | - | 0.96 | -0.02075 | 0.00652 | 245 | -3.182 | .017 | -0.6364 |
| | - | 0.98 | -0.02275 | 0.00652 | 245 | -3.489 | .006 | -0.6977 |
| | - | 1.00 | -0.50925 | 0.00652 | 245 | -78.094 | <.001 | -15.6189 |
| 0.94 | - | 0.96 | -7.50e−4 | 0.00652 | 245 | -0.115 | 1.000 | -0.0230 |
| | - | 0.98 | -0.00275 | 0.00652 | 245 | -0.422 | 1.000 | -0.0843 |
| | - | 1.00 | -0.48925 | 0.00652 | 245 | -75.027 | <.001 | -15.0055 |
| 0.96 | - | 0.98 | -0.00200 | 0.00652 | 245 | -0.307 | 1.000 | -0.0613 |
| | - | 1.00 | -0.48850 | 0.00652 | 245 | -74.912 | <.001 | -14.9824 |
| 0.98 | - | 1.00 | -0.48650 | 0.00652 | 245 | -74.606 | <.001 | -14.9211 |

Note. Comparisons based on estimated marginal means.

**Table S6**

Results of the post hoc tests for the decision accuracy when manipulating transition precision (under high reward value).

| Transition Precision | | | | | | | | |
|---|---|---|---|---|---|---|---|---|
| condition | | condition | Mean difference | Standard Error | dof | t | $p_{bonferroni}$ | Cohen's d |
| 0.92 | - | 0.94 | -0.0120 | 0.00534 | 245 | -2.25 | .256 | -0.449 |
| | - | 0.96 | -0.0285 | 0.00534 | 245 | -5.33 | <.001 | -1.067 |
| | - | 0.98 | -0.0667 | 0.00534 | 245 | -12.49 | <.001 | -2.498 |
| | - | 1.00 | -0.0780 | 0.00534 | 245 | -14.60 | <.001 | -2.920 |
| 0.94 | - | 0.96 | -0.0165 | 0.00534 | 245 | -3.09 | .022 | -0.618 |
| | - | 0.98 | -0.0547 | 0.00534 | 245 | -10.25 | <.001 | -2.049 |
| | - | 1.00 | -0.0660 | 0.00534 | 245 | -12.35 | <.001 | -2.470 |
| 0.96 | - | 0.98 | -0.0382 | 0.00534 | 245 | -7.16 | <.001 | -1.432 |
| | - | 1.00 | -0.0495 | 0.00534 | 245 | -9.26 | <.001 | -1.853 |
| 0.98 | - | 1.00 | -0.0113 | 0.00534 | 245 | -2.11 | .363 | -0.421 |

Note. Comparisons based on estimated marginal means.

**Table S7**

Results of the post hoc tests for the proportion of reward collected when manipulating transition precision (under high reward value).

| Transition Precision | | | | | | | | |
|---|---|---|---|---|---|---|---|---|
| condition | | condition | Mean difference | Standard Error | dof | t | $p_{bonferroni}$ | Cohen's d |
| 0.92 | - | 0.94 | 0.0199 | 0.00735 | 245 | 2.71 | .072 | 0.542 |
| | - | 0.96 | 0.0412 | 0.00735 | 245 | 5.61 | <.001 | 1.123 |
| | - | 0.98 | -0.5631 | 0.00735 | 245 | -76.62 | <.001 | -15.324 |
| | - | 1.00 | -0.5812 | 0.00735 | 245 | -79.08 | <.001 | -15.816 |
| 0.94 | - | 0.96 | 0.0213 | 0.00735 | 245 | 2.90 | .040 | 0.581 |
| | - | 0.98 | -0.5830 | 0.00735 | 245 | -79.33 | <.001 | -15.866 |
| | - | 1.00 | -0.6011 | 0.00735 | 245 | -81.79 | <.001 | -16.358 |
| 0.96 | - | 0.98 | -0.6043 | 0.00735 | 245 | -82.23 | <.001 | -16.446 |
| | - | 1.00 | -0.6224 | 0.00735 | 245 | -84.69 | <.001 | -16.938 |
| 0.98 | - | 1.00 | -0.0181 | 0.00735 | 245 | -2.46 | .146 | -0.492 |

Note. Comparisons based on estimated marginal means.

**Table S8**

Results of the post hoc tests for the decision accuracy when manipulating sensory precision (under constant reward value).

| Precision in Likelihood Mapping | | | | | | | | |
|---|---|---|---|---|---|---|---|---|
| condition | | condition | Mean difference | Standard Error | dof | t | $p_{bonferroni}$ | Cohen's d |
| 0.92 | - | 0.94 | -0.00500 | 0.00534 | 245 | -0.937 | 1.000 | -0.187 |
| | - | 0.96 | -0.03250 | 0.00534 | 245 | -6.091 | <.001 | -1.218 |
| | - | 0.98 | -0.05300 | 0.00534 | 245 | -9.932 | <.001 | -1.986 |
| | - | 1.00 | -0.07525 | 0.00534 | 245 | -14.102 | <.001 | -2.820 |
| 0.94 | - | 0.96 | -0.02750 | 0.00534 | 245 | -5.154 | <.001 | -1.031 |
| | - | 0.98 | -0.04800 | 0.00534 | 245 | -8.995 | <.001 | -1.799 |
| | - | 1.00 | -0.07025 | 0.00534 | 245 | -13.165 | <.001 | -2.633 |
| 0.96 | - | 0.98 | -0.02050 | 0.00534 | 245 | -3.842 | .002 | -0.768 |
| | - | 1.00 | -0.04275 | 0.00534 | 245 | -8.012 | <.001 | -1.602 |
| 0.98 | - | 1.00 | -0.02225 | 0.00534 | 245 | -4.170 | <.001 | -0.834 |

Note. Comparisons based on estimated marginal means.

**Table S9**

Results of the post hoc tests for the proportion of reward collected when manipulating sensory precision (under constant reward value).

| Precision in Likelihood Mapping | | | | | | | | |
|---|---|---|---|---|---|---|---|---|
| condition | | condition | Mean difference | Standard Error | dof | t | $p_{bonferroni}$ | Cohen's d |
| 0.92 | - | 0.94 | -0.0100 | 0.0107 | 245 | -0.937 | 1.000 | -0.187 |
| | - | 0.96 | -0.0650 | 0.0107 | 245 | -6.091 | <.001 | -1.218 |
| | - | 0.98 | -0.1060 | 0.0107 | 245 | -9.932 | <.001 | -1.986 |
| | - | 1.00 | -0.1505 | 0.0107 | 245 | -14.102 | <.001 | -2.820 |
| 0.94 | - | 0.96 | -0.0550 | 0.0107 | 245 | -5.154 | <.001 | -1.031 |
| | - | 0.98 | -0.0960 | 0.0107 | 245 | -8.995 | <.001 | -1.799 |
| | - | 1.00 | -0.1405 | 0.0107 | 245 | -13.165 | <.001 | -2.633 |
| 0.96 | - | 0.98 | -0.0410 | 0.0107 | 245 | -3.842 | .002 | -0.768 |
| | - | 1.00 | -0.0855 | 0.0107 | 245 | -8.012 | <.001 | -1.602 |
| 0.98 | - | 1.00 | -0.0445 | 0.0107 | 245 | -4.170 | <.001 | -0.834 |

Note. Comparisons based on estimated marginal means.

**Table S10**

Results of the post hoc tests for the decision accuracy when manipulating sensory precision (under medium reward value).

| Precision in Likelihood Mapping | | | | | | | | |
|---|---|---|---|---|---|---|---|---|
| condition | | condition | Mean difference | Standard Error | dof | t | $p_{bonferroni}$ | Cohen's d |
| 0.92 | - | 0.94 | -0.0157 | 0.00524 | 245 | -3.01 | .029 | -0.601 |
| | - | 0.96 | -0.0355 | 0.00524 | 245 | -6.78 | <.001 | -1.355 |
| | - | 0.98 | -0.0582 | 0.00524 | 245 | -11.12 | <.001 | -2.223 |
| | - | 1.00 | -0.0712 | 0.00524 | 245 | -13.60 | <.001 | -2.720 |
| 0.94 | - | 0.96 | -0.0198 | 0.00524 | 245 | -3.77 | .002 | -0.754 |
| | - | 0.98 | -0.0425 | 0.00524 | 245 | -8.11 | <.001 | -1.622 |
| | - | 1.00 | -0.0555 | 0.00524 | 245 | -10.59 | <.001 | -2.118 |
| 0.96 | - | 0.98 | -0.0227 | 0.00524 | 245 | -4.34 | <.001 | -0.868 |
| | - | 1.00 | -0.0358 | 0.00524 | 245 | -6.82 | <.001 | -1.365 |
| 0.98 | - | 1.00 | -0.0130 | 0.00524 | 245 | -2.48 | .138 | -0.496 |

Note. Comparisons based on estimated marginal means.

**Table S11**

Results of the post hoc tests for the proportion of reward collected when manipulating sensory precision (under medium reward value).

| Precision in Likelihood Mapping | | | | | | | | |
|---|---|---|---|---|---|---|---|---|
| condition | | condition | Mean difference | Standard Error | dof | t | $p_{bonferroni}$ | Cohen's d |
| 0.92 | - | 0.94 | -0.0157 | 0.00545 | 245 | -2.89 | .042 | -0.578 |
| | - | 0.96 | -0.0355 | 0.00545 | 245 | -6.51 | <.001 | -1.302 |
| | - | 0.98 | -0.0582 | 0.00545 | 245 | -10.68 | <.001 | -2.137 |
| | - | 1.00 | -0.5618 | 0.00545 | 245 | -103.04 | <.001 | -20.607 |
| 0.94 | - | 0.96 | -0.0198 | 0.00545 | 245 | -3.62 | .004 | -0.725 |
| | - | 0.98 | -0.0425 | 0.00545 | 245 | -7.80 | <.001 | -1.559 |
| | - | 1.00 | -0.5460 | 0.00545 | 245 | -100.15 | <.001 | -20.029 |
| 0.96 | - | 0.98 | -0.0227 | 0.00545 | 245 | -4.17 | <.001 | -0.835 |
| | - | 1.00 | -0.5263 | 0.00545 | 245 | -96.52 | <.001 | -19.305 |
| 0.98 | - | 1.00 | -0.5035 | 0.00545 | 245 | -92.35 | <.001 | -18.470 |

Note. Comparisons based on estimated marginal means.

**Table S12**

Results of the post hoc tests for the decision accuracy when manipulating sensory precision (under high reward value).

| Precision in Likelihood Mapping | | | | | | | | |
|---|---|---|---|---|---|---|---|---|
| condition | | condition | Mean difference | Standard Error | dof | t | $p_{bonferroni}$ | Cohen's d |
| 0.92 | - | 0.94 | -0.00950 | 0.00749 | 245 | -1.27 | 1.000 | -0.254 |
| | - | 0.96 | -0.02400 | 0.00749 | 245 | -3.21 | .015 | -0.641 |
| | - | 0.98 | -0.26350 | 0.00749 | 245 | -35.19 | <.001 | -7.039 |
| | - | 1.00 | -0.27850 | 0.00749 | 245 | -37.20 | <.001 | -7.440 |
| 0.94 | - | 0.96 | -0.01450 | 0.00749 | 245 | -1.94 | .539 | -0.387 |
| | - | 0.98 | -0.25400 | 0.00749 | 245 | -33.93 | <.001 | -6.785 |
| | - | 1.00 | -0.26900 | 0.00749 | 245 | -35.93 | <.001 | -7.186 |
| 0.96 | - | 0.98 | -0.23950 | 0.00749 | 245 | -31.99 | <.001 | -6.398 |
| | - | 1.00 | -0.25450 | 0.00749 | 245 | -33.99 | <.001 | -6.798 |
| 0.98 | - | 1.00 | -0.01500 | 0.00749 | 245 | -2.00 | .462 | -0.401 |

Note. Comparisons based on estimated marginal means.

**Table S13**

Results of the post hoc tests for the proportion of reward collected when manipulating sensory precision (under high reward value).

| Precision in Likelihood Mapping | | | | | | | | |
|---|---|---|---|---|---|---|---|---|
| condition | | condition | Mean difference | Standard Error | dof | t | $p_{bonferroni}$ | Cohen's d |
| 0.92 | - | 0.94 | -0.00967 | 0.0131 | 245 | -0.736 | 1.000 | -0.147 |
| | - | 0.96 | -0.02600 | 0.0131 | 245 | -1.979 | .490 | -0.396 |
| | - | 0.98 | -0.49033 | 0.0131 | 245 | -37.314 | <.001 | -7.463 |
| | - | 1.00 | -0.51000 | 0.0131 | 245 | -38.811 | <.001 | -7.762 |
| 0.94 | - | 0.96 | -0.01633 | 0.0131 | 245 | -1.243 | 1.000 | -0.249 |
| | - | 0.98 | -0.48067 | 0.0131 | 245 | -36.578 | <.001 | -7.316 |
| | - | 1.00 | -0.50033 | 0.0131 | 245 | -38.075 | <.001 | -7.615 |
| 0.96 | - | 0.98 | -0.46433 | 0.0131 | 245 | -35.336 | <.001 | -7.067 |
| | - | 1.00 | -0.48400 | 0.0131 | 245 | -36.832 | <.001 | -7.366 |
| 0.98 | - | 1.00 | -0.01967 | 0.0131 | 245 | -1.497 | 1.000 | -0.299 |

Note. Comparisons based on estimated marginal means.
